## Supplementary Figures for "Isolation and characterisation of bacteriophages with activity against invasive non-typhoidal *Salmonella* causing bloodstream infection in Malawi"

**Figure S1:** Plaque morphology used for defining host range (Figure 1). Plaques were designated “defined” where there was distinct zone of clearance, otherwise plaques were classified as “smaller” when presenting smaller size, or “indistinct” when presented undefined edges.

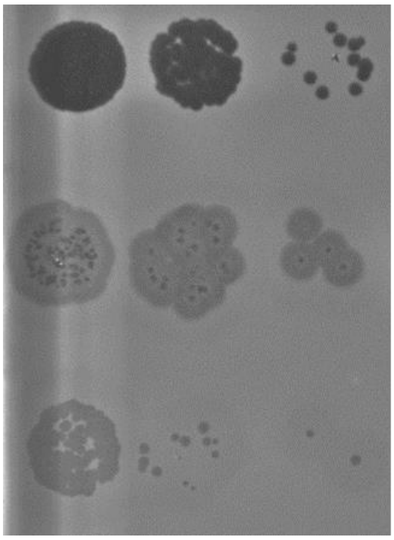

Defined plaques

Indistinct plaques

Smaller plaques

**Figure S2:** Example of Random Amplified Polymorphic DNA (RAPD) profiles. Distinct individual plaques were subjected to RAPD-PCR using P1, P2, OPL5 and RAPD5 primers (Supplementary Table S1), visualised by 0.8% agarose gel electrophoresis at 100 V for 60 min. L indicates 1 kb ladder (Bioline, H1-819101A) and -ve indicates the negative control that lacked template. Plaques 1-15 are examples of different patterns obtained with primers P1 (A) or RAPD5 (B), and plaques 16-32 are examples of different patterns obtained with primers P2 and OPL5.

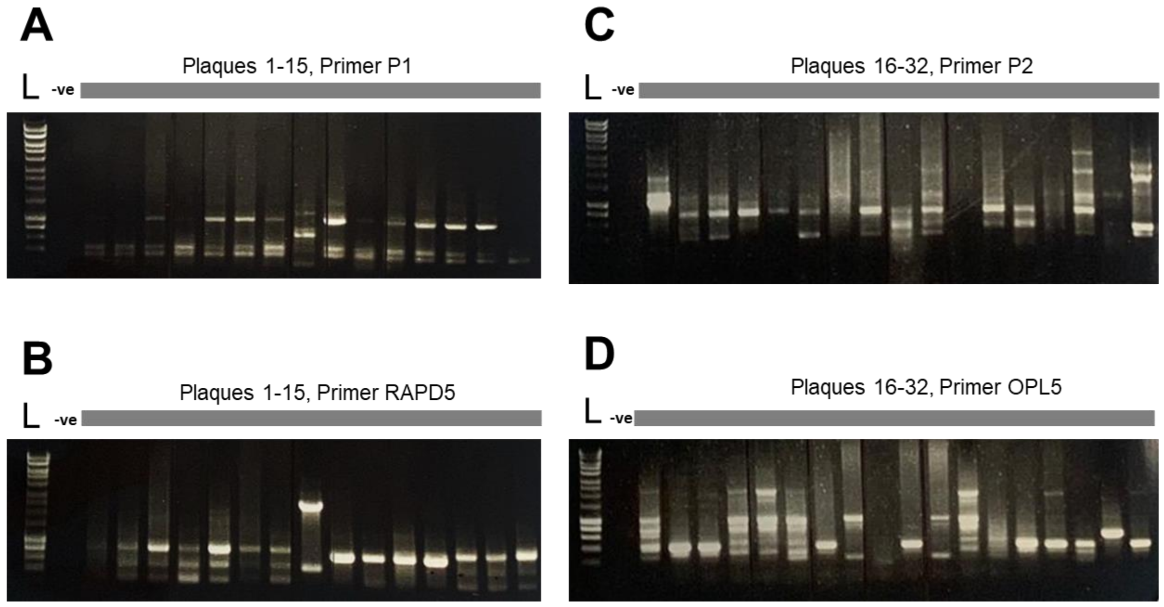

**Figure S3:** Comparative genomic analysis of phages ER25 and P22. The upper panel includes nucleotides 1 to 19,950 of P22, and the bottom panel includes nucleotides 19,951 to the end of P22. Arrows indicate annotated genes and the 49 SNPs and indels identified are indicated in red.

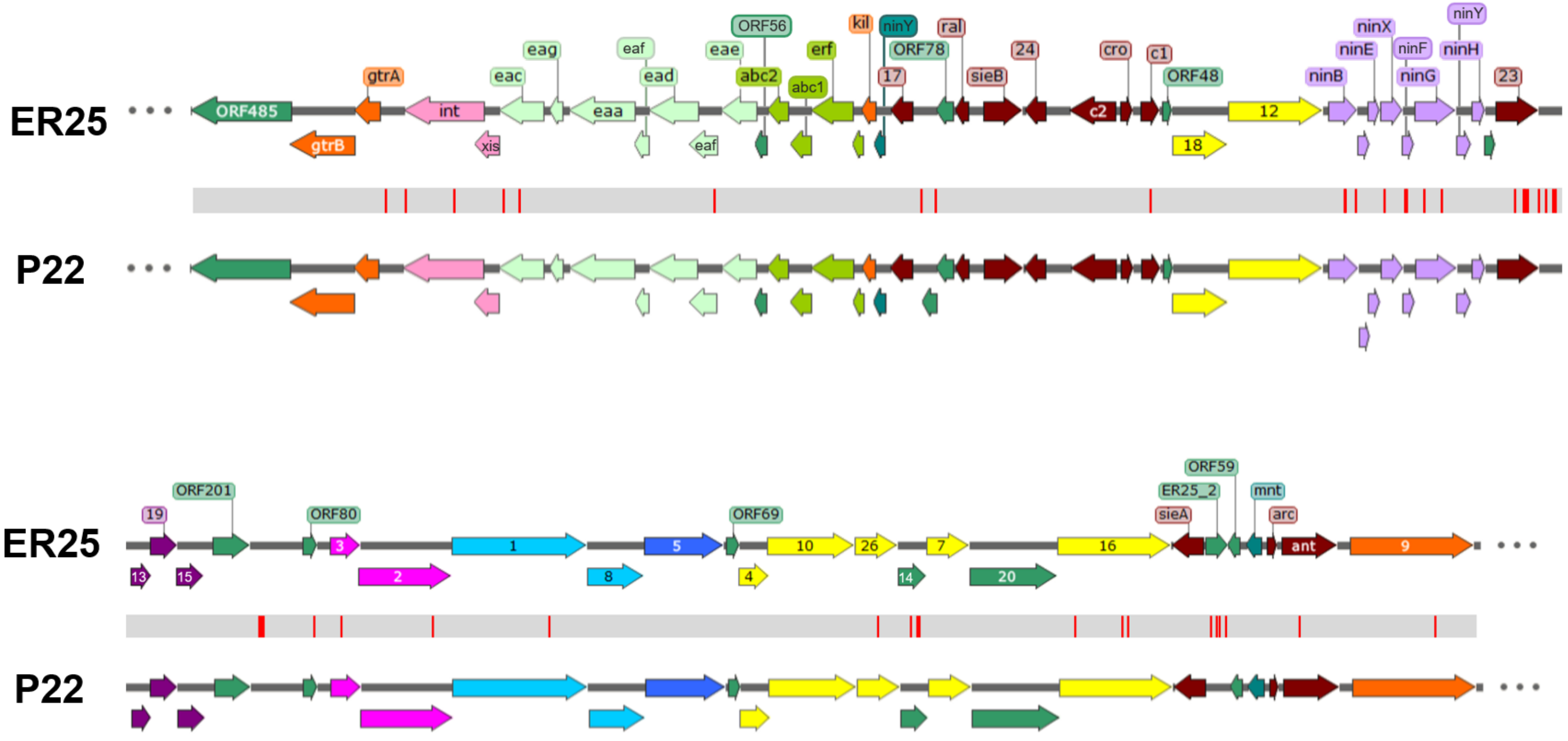
